# Divergent behavioral responses in protracted opioid withdrawal in male and female C57BL/6J mice

**DOI:** 10.1101/689521

**Authors:** Isabel M. Bravo, Brennon R. Luster, Meghan E. Flanigan, Patric J. Perez, Elizabeth S. Cogan, Karl T. Schmidt, Zoe A. McElligott

## Abstract

Persons suffering from opioid use disorder (OUD) experience long-lasting dysphoric symptoms well into extended periods of withdrawal. This protracted withdrawal syndrome is notably characterized by heightened anxiety. Here we investigate if an exacerbated withdrawal model of acute morphine dependence results in lasting behavioral adaptation 6 weeks into forced abstinence. We found that our exacerbated morphine withdrawal paradigm produced distinct impairments in elevated-plus maze, open field, and social interaction tests in male and female mice. These findings will be relevant for future investigation examining the neural mechanisms underlying these behaviors, and will aid in uncovering physiological sex differences in response to opioid withdrawal.

## Introduction

Opioid use disorder (OUD) is a chronic relapsing condition in which individuals remain vulnerable to drug craving, anxiety, and relapse even after prolonged periods of abstinence (Sinha, 2008; Kreek *et al.*, 2010; Kreek, 2011). The current opioid epidemic is largely driven by an over-prescription of legal opioids (Center for Behavioral Health Statistics and Quality, 2016). Despite the brevity of initial exposure, individuals subsequently experience varying degrees of a withdrawal syndrome (Heishman *et al.*, 1989, 1990; Kirby *et al.*, 1990; Kirby & Stitzer, 1993) that leads to long-lasting behavioral and biological changes which increase propensity for future drug use. There is strong evidence that such plasticity differs between males and females, demonstrated by differences observed in the analgesic effects of opioids (Lee & Ho, 2013), response to drug-related cues (Yu *et al.*, 2007) and responsivity and progression of drug-taking, suggesting that it is crucial to investigate how opioids may differentially modulate behavior in males and females.

Our lab has adapted a three-day morphine dependence paradigm (Luster *et al.*, 2019), which was used in this study to determine the effects of repeated bouts of morphine withdrawal. As observed in clinical settings and other established animal models, this model produces robust sensitization of physical withdrawal symptoms from morphine across the three-day period (Azorlosa & Stitzer, 1990; Azolosa *et al.*, 1994; Schulteis *et al.*, 1998, 1999), and acute anxiety-like behavior in morphine-treated animals (McElligott et al 2013). It was unclear, however, if this paradigm, designed to examine acute withdrawal effects, would have lasting effects on mouse behavior after several weeks of abstinence, and whether there would be sexually dimorphic effects. Since addiction is a chronic relapsing disorder, it is crucial to study the long-term changes that occur as a result of initial drug intake. Thus, a valid model of morphine dependence should demonstrate long-term changes that can contribute to relapse. While it is challenging to develop a model of opioid dependence and withdrawal in mice with face validity, we propose that our model produces behavioral and physiological effects that parallel clinical observations.

The current set of experiments were conducted to determine if (1) there would be lasting behavioral consequences to this exacerbated withdrawal paradigm, and (2) whether male and female mice would exhibit different anxiety profiles after a period of abstinence. Here we demonstrate that both male and female C57BL/6J mice have altered behavior following 6 weeks of protracted morphine withdrawal, suggesting long-term behavioral plasticity. Moreover, we observed striking sex differences in the assays we probed. Finally, previously withdrawn male and female mice both showed elevated responsivity to subsequent morphine challenge, however at different doses. Our findings demonstrate that this model produces protracted withdrawal symptoms that are reminiscent of clinical reports.

## Methods

### Subjects

All procedures were approved by the University of North Carolina Institutional Animal Use and Care Committee. Male and female C57BL/6J mice at least 10 weeks of age were used in all experiments and maintained on a normal 12:12 light dark cycle, in both single and group housed conditions. All animals had food and water *ad libitum* for the duration of the study. Animals were treated for 3 days with the following paradigm, as described previous (Luster *et al.*, 2019). Briefly, Saline Naloxone (SN) animals were injected with (0.9%) sterile saline (0.1 ml/10g) and 2 hours later with 1 mg/kg naloxone (in sterile saline 0.1 ml/10g). Morphine Naloxone (MN) animals were injected with 10 mg/kg morphine (in sterile saline 0.1 ml/10 g) and 2 hours later with 1 mg/kg naloxone (in sterile saline 0.1 ml/10 g). Injections were delivered subcutaneously. All animals were observed for withdrawal behavior, and all morphine animals exhibited classical withdrawal signs (e.g., escape jumps, paw tremor, teeth chattering) as we have previously reported for male and female C57BL/6J mice (Luster *et al.*, 2019). Following the third day of the paradigm, animals were undisturbed in the animal facility with the exception of routine cage changes for 6 weeks before testing.

### Anxiety-like assays

#### Elevated Plus Maze

The elevated plus maze (EPM) consists of two open arms measuring 77 cm × 77 cm and two closed arms of the same length(77 cm × 77 cm). Arms of the EPM meet at a common central platform, and the maze is elevated 74 cm above the ground. Light was measured to be 150 lux in the center of the maze. Mice were placed on the central platform and an overhead camera was used to record a 5-minute testing session. Videos were analyzed using Ethovision software (Noldus, Netherlands).

#### Open Field

Mice were placed in an open field. Locomotor activity and position within the open field was measured for 60 minutes. The open field consisted of a white plastic box measuring 50 cm × 50 cm within a sound attenuated chamber. Light in the chamber was measured to be 40 lux. Videos were analyzed using Ethovision software (Noldus, Netherlands).

#### Free Social Interaction Testing

Singly housed experimental mice (1 week prior to testing) and group housed social target mice were habituated to the testing arena for 10 minutes one day before testing. The testing arena consisted of a clear plastic mouse cage with dimensions 33.6 cm long × 18.1 cm wide × 13.8 cm deep (no bedding present), 45 lux. On test day, experimental mice and novel, drug-naïve, age- and sex-matched social target mice were placed into the testing arena and behavior was recorded for 10 minutes using an overhead camera. To distinguish social target mice from experimental mice, the backs of social target mice were shaved. The duration of time that experimental mice spent actively engaging in social behavior was measured by a blind observer (i.e. the experimental mouse must have been oriented towards the social target mouse). Time that the experimental mouse spent passively engaging in social behavior was not counted towards this duration (i.e. if the social target mouse was engaging in ano-genital sniffing of the experimental mouse).

#### *Morphine* C*hallenge Doses*

Six weeks post withdrawal, mice were probed for long lasting behavioral adaptations with increasing challenge doses of morphine. On test days mice were placed in the behavioral testing room for a habituation period of 120 minutes. After the habituation period mice received a subcutaneous injection (saline, or morphine: 1, 5, 10 mg/kg) and were placed individually in an open field inside a locomotor activity chamber (Accuscan Instruments). The open field consisted of a 40.6 cm square clear plexiglass box within a sound attenuated cubical. Light in the cubicle was measured as 40 Lux. Mice were allowed to explore the open field for 150 minutes while their locomotor activity was recorded. At the end of the testing period, mice were returned to home cages. All mice received saline, and 3 different doses of morphine (1, 5, 10 mg/kg) on separate testing days with at least two days between each testing day.

### Statistics

All data were analyzed using Graph Pad Prism (versions 6, 7, and 8). Data is reported as mean ± SEM. Comparisons were made with either unpaired Student’s t-test, or repeated measures 2-way ANOVAS. One SN male was removed from the data set as a statistical outlier (Grubb’s test.)

## Results

### Baseline anxiety behavior in protracted withdrawal

To model opioid dependence in C57BL/6J mice, we used a 3-day paradigm of exacerbated withdrawal (**Fig1**). We recently demonstrated that this paradigm enhances the development of withdrawal syndrome across the 3 days in both male and female mice, and alters inhibitory transmission in the BNST (Luster *et al.*, 2019). Interestingly, however, we found that both the manifestation of somatic withdrawal responses and BNST inhibitory transmission were impacted in sexually dimorphic ways (Luster *et al.*, 2019). While this paradigm was developed as an acute dependence model (Schulteis *et al.*, 1998, 1999; McElligott *et al.*, 2013), we hypothesized that there may be lasting changes in animal behavior in protracted abstinence. Therefore, we investigated behavior in male and female animals 6 weeks following the last morphine/saline and naloxone pairing. Interestingly, we found that both male and female mice who had experienced morphine withdrawal showed lasting, albeit divergent, behavioral adaptations.

**Figure 1.**
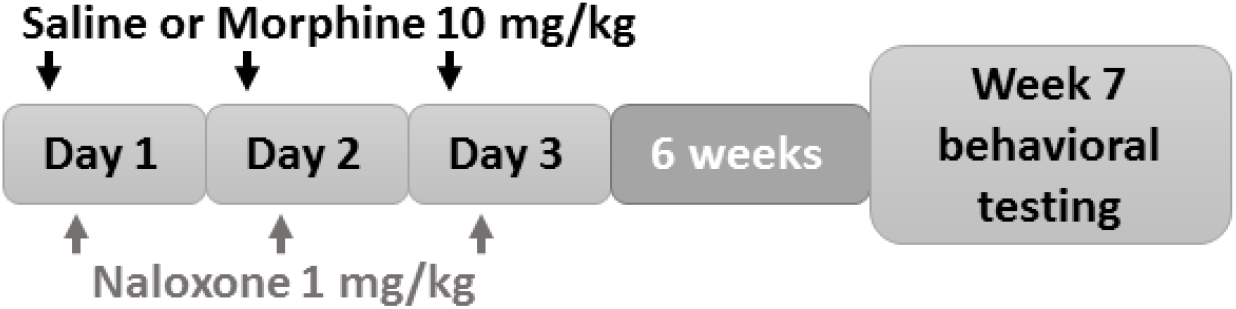
Schematic of morphine withdrawal model. Equal volume (0.1 mL/10 g) saline or 10mg/kg morphine subcutaneously (s.c.) was injected and 1 mg/kg naloxone s.c. was injected 2 hours later. This paradigm was repeated for a total of 3 days. Subsequently animals experienced forced abstinence for 6 weeks, prior to behavioral testing.

Six weeks following the final day of the paradigm, we assessed the animals’ behavior at baseline in two assays that put the mouse’s proclivity to explore a novel environment in conflict with a desire to stay safe. First, we evaluated the animals in the elevated plus maze (EPM). All animals were placed into the center of the maze, and allowed to explore the maze for 5 minutes. Compared to SN male mice, MN male mice spent a significantly longer period of time in the open arms of the maze (SNM: 14.9 ± 2.8%, n = 10; MNM: 28.6 ± 5.8% n = 9; % total time, p<0.05; **Fig2A**) of total time, and a significantly shorter period of time in the closed arms (SNM: 70.5 ± 3.2%, MNM: 53.9 ± 7.1%, p<0.05; **Fig2C**). Additionally, MN male mice had a significantly decreased latency to enter the open arms as compared to their SN controls (SNM: 14.0 ± 3.9 s; MNM: 1.4 ± 1.1 s, p<0.01; **Fig2B**). We did not observe differences in center time (SNM: 15.3 ± 1.7%: MNM: 18.2 ± 2.3%; **Fig2D**), total distance traveled (SNM: 1402 ± 101.7 cm; MNM: 1224 ± 93.2 cm; **Fig2F**), or velocity (SNM: 4.7 ± 0.3 cm/s; MNM: 4.1 ± 0.3 cm/s; **Fig2E**). In contrast, we did not observe any significant differences between female MN or SN mice on any metrics: % time in the open arm (SNF: 18.4 ± 2.1%, n = 8; MNF: 18.6 ± 3.9%, n = 7; **Fig3A**), % time in the closed arm (SNF: 61.5 ± 1.6%, MNF: 57.7 ± 3.3%; **Fig3C**), % time in center (SNF: 19.1 ± 2.1%, MNF: 21.7 ± 3.0%; **Fig3D**); latency to enter the open arm (SNF: 5.0 ± 4.0 s; MNF: 6.5 ± 4.1 s; **Fig3B**), distance traveled (SNF: 1439 ± 102.3 cm; MNF: 1485 ± 75.0 cm; **Fig3F**); velocity (SNF: 4.5 ± 0.3 cm/s; MNF: 4.4 ± 0.3 cm/s; **Fig3E**).

**Figure 2.**
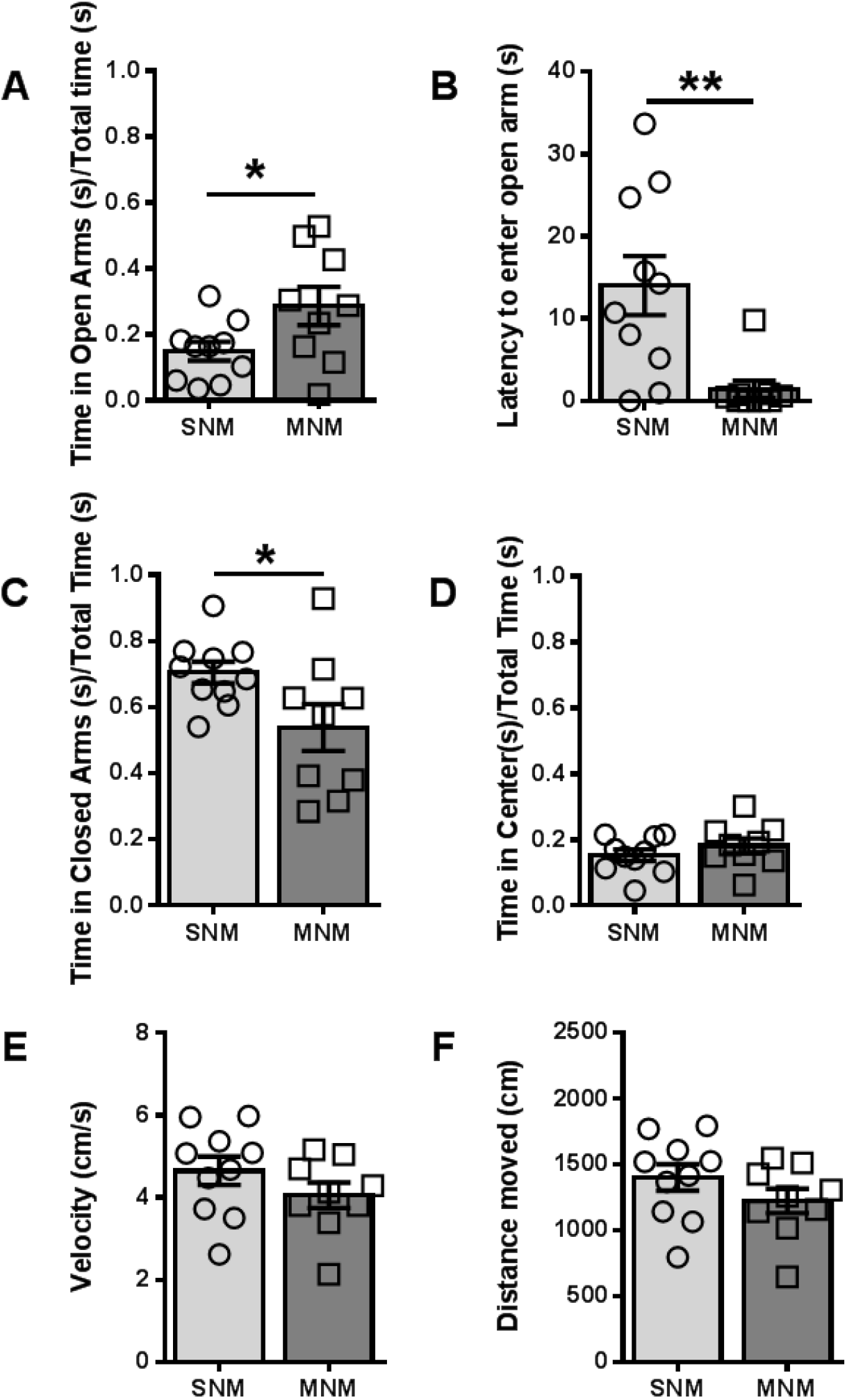
Male mice show altered exploration of the elevated plus maze in protracted withdrawal. A. Ratio of time spent in the open arms over total time. B. Latency (s) for mice to enter into the open arm. C. Ratio of time spent in the closed arms over total time. D. Ratio of time spent in the center of the EPM over total time. E. Velocity of mouse in the EPM (cm/s). F. Distance traveled in the EPM (cm). * p<0.05, ** p<0.01

**Figure 3.**
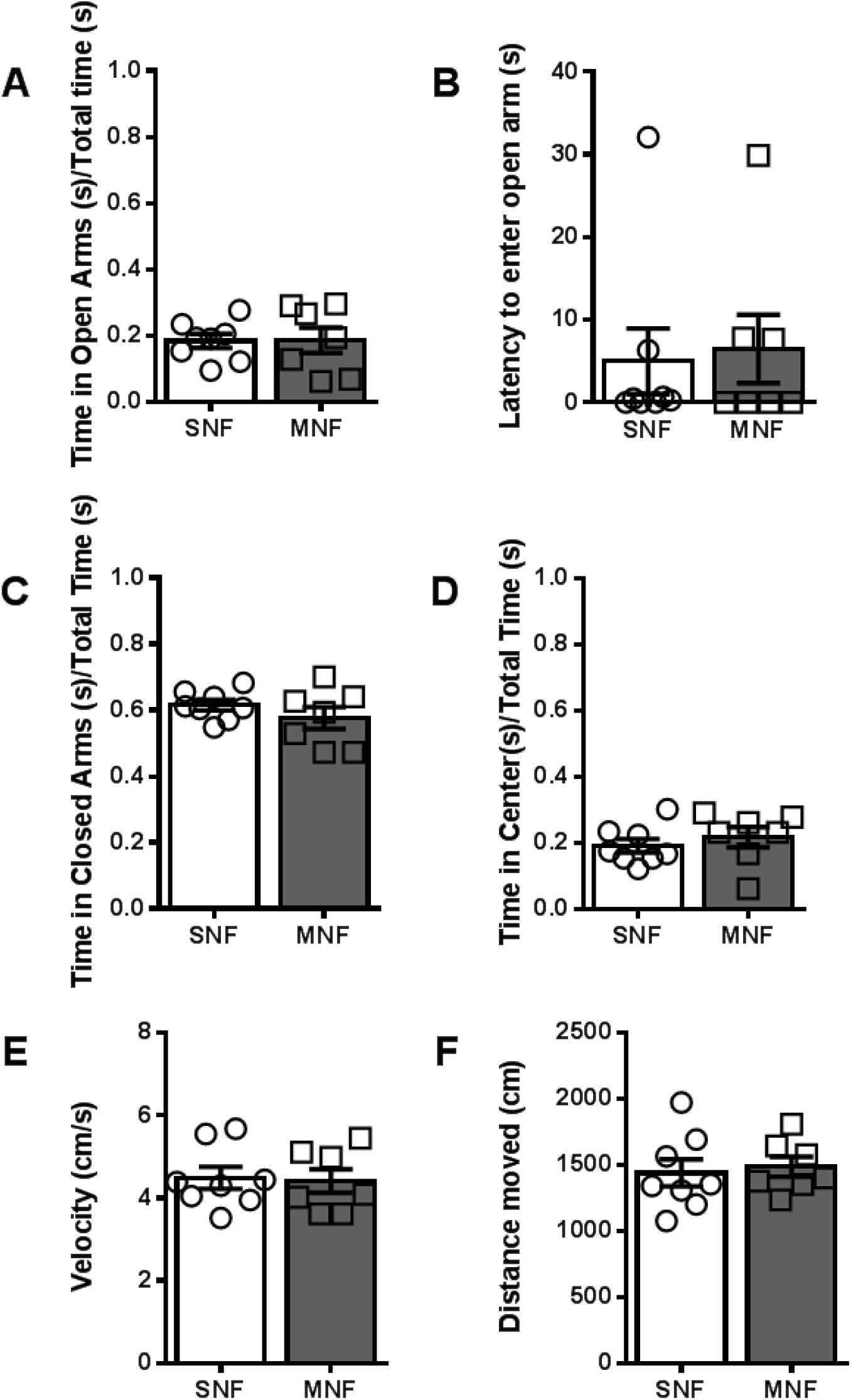
Female mice do not exhibit differences in the elevated plus maze in protracted withdrawal. A. Ratio of time spent in the open arms over total time. B. Latency (s) for mice to enter into the open arm. C. Ratio of time spent in the closed arms over total time. D. Ratio of time spent in the center of the EPM over total time. E. Velocity of mouse in the EPM (cm/s). F. Distance traveled in the EPM (cm).

We next evaluated the animals in a 60 minute open field (OF) session. We found that male mice did not exhibit a difference in center (SNM: 8.0 ± 0.7%, n = 11; MNM: 7.5 ± 1.1%, n = 13; **Fig4A,D**), surround (SNM: 92.0 ± 0.7%; MNM: 92.5 ± 1.1%, **Fig4B,E**), or corner time (SNM: 56.0 ± 1.7%; MNM: 59.4 ± 2.9%; **Fig4C,F**). Previous morphine withdrawal significantly increased the latency to enter the corners (SNM: 0.8 ± 0.3 s; MNM: 3.6 ± 1.1 s; p<0.05; **Fig4G**), consistent with the behavior of male mice in the EPM. In contrast to the males, MN female mice exhibited several behavioral differences in the open field as compared to their SN controls. Female mice in protracted withdrawal had spent less time in the center (SNF: 8.0 ± 0.5%, n = 8; MNF: 6.5 ± 0.3%, n = 7; p<0.05; **Fig5A,D**), enhanced time in the surround (SNF: 92.0 ± 0.5%; MNF: 93.5 ± 3.0%; p<0.05; **Fig5B,E**), and enhanced time in the corners (SNF: 47.6 ± 1.3%; MNF: 52.0 ± 1.2%; p<0.05, **Fig5C,F**).

**Figure 4.**
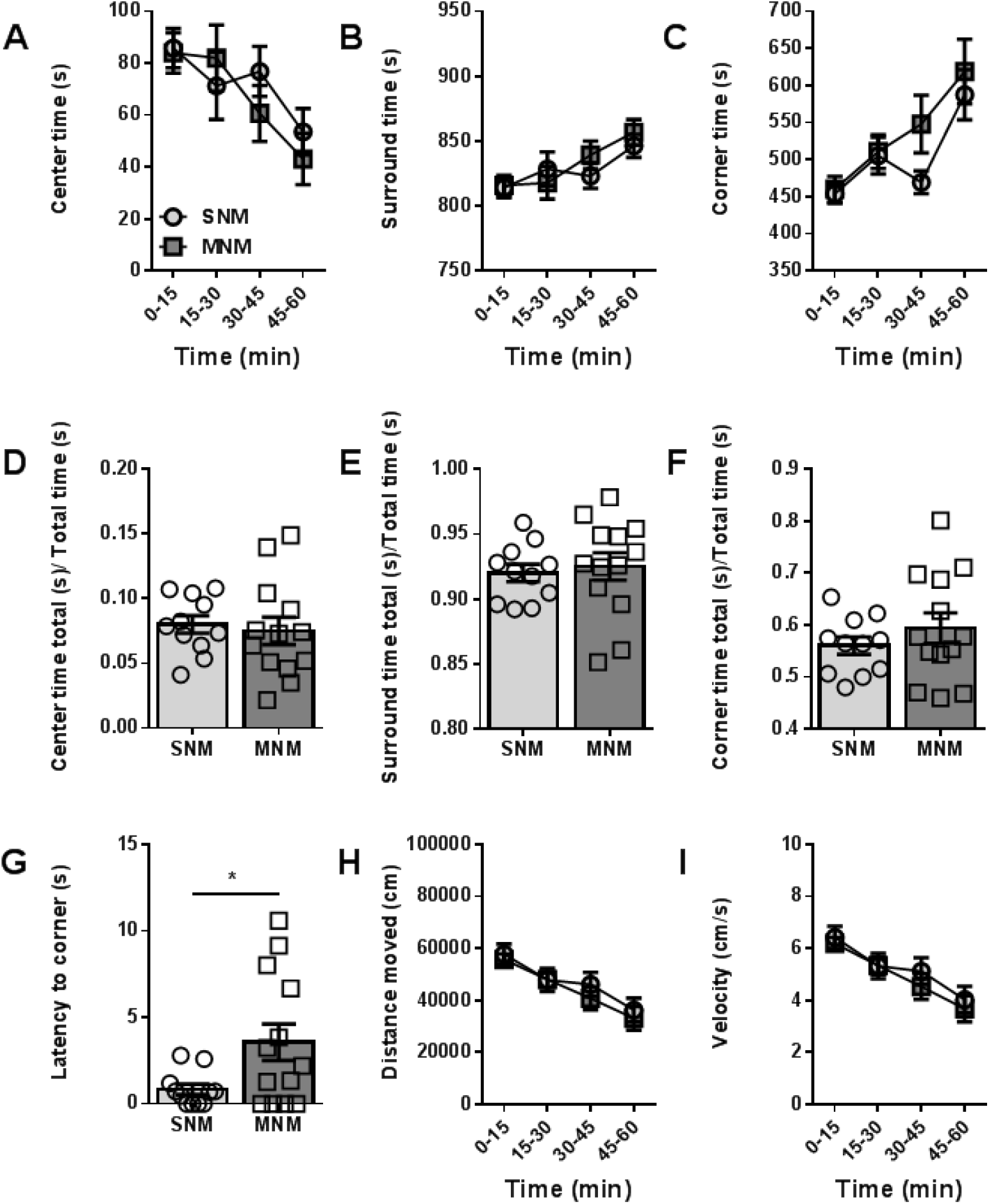
Protracted withdrawal alters latency to enter corners of an open field in male mice. A. Ratio of time spent in the center of the open field over total time in 15-minute time bins. B. Ratio of time spent in the surround of the open field over total time in 15 min time bins. C. Ratio of time spent in the corners of the open field over total time in 15-minute time bins. D. Ratio of time spent in the center over total time. E. Ratio of time spent in the surround over total time. F. Ratio of time spent in the corners over total time. G. Latency to enter the corner of the open field (s). H. Distance moved in the open field (cm). I. Velocity in the open field (cm/s). * p>0.05

**Figure 5.**
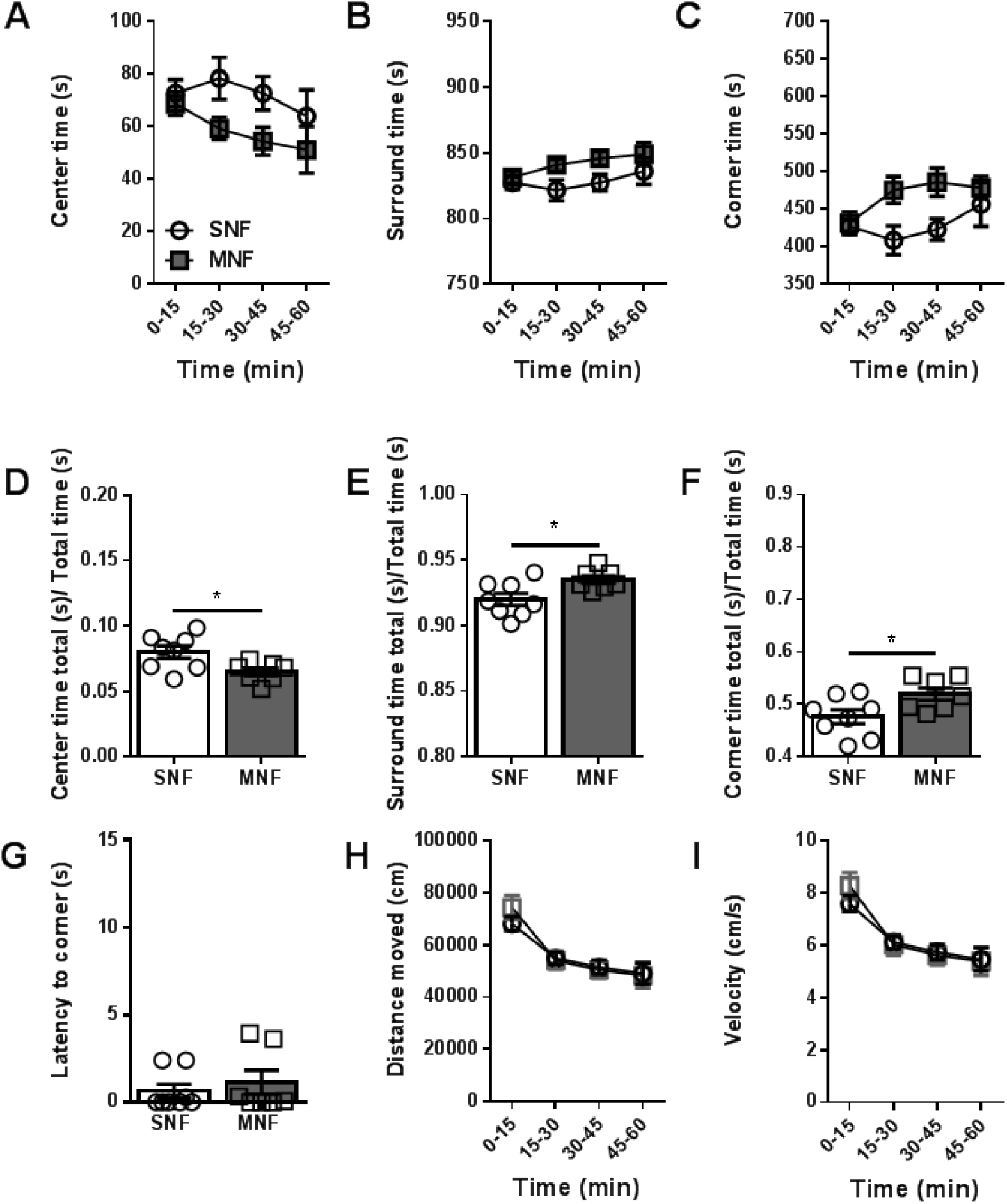
Protracted withdrawal alters time spent in center and corner of the open field in female mice. A. Ratio of time spent in the center of the open field over total time in 15-minute time bins. B. Ratio of time spent in the surround of the open field over total time in 15-minute time bins. C. Ratio of time spent in the corners of the open field over total time in 15 min time bins. D. Ratio of time spent in the center over total time. E. Ratio of time spent in the surround over total time. F. Ratio of time spent in the corners over total time. G. Latency to enter the corner of the open field (s). H. Distance moved in the open field (cm). I. Velocity in the open field (cm/s). * p>0.05

Because of the behavioral differences we observed in the EPM and open field between male and female mice, we next examined how mice would respond to a more ethologically-relevant challenge, free social interaction. Social anxiety disorder is highly co-morbid in clinical populations abusing opioids and other substances (Applebaum *et al.*, 2010; Shand *et al.*, 2010). We hypothesized that mice in protracted withdrawal would exhibit reduced interaction with a novel mouse. Six weeks after the last precipitated withdrawal, mice were placed in a standard mouse cage along with a naive, age and sex matched target mouse. We scored the duration of time that the experimental mouse spent actively interacting with the target mouse over the course of 10 minutes. We defined active social interaction to be any time the experimental mouse was in close contact with and oriented towards the target mouse. Interestingly, we found that male mice who had previously experienced morphine withdrawal spent significantly less time interacting with the conspecific versus control mice (SNM: 23.3 ± 3.7%; MNM: 13.1 ± 4.0%; n = 5 both groups; p<0.05; **Fig 6A**). In contrast, we did not observe differences in the female mice (SNF: 15.0 ± 7.2%; MNF: 13.7 ± 3.2%; n = 5 both groups; **Fig 6B**).

**Figure 6.**
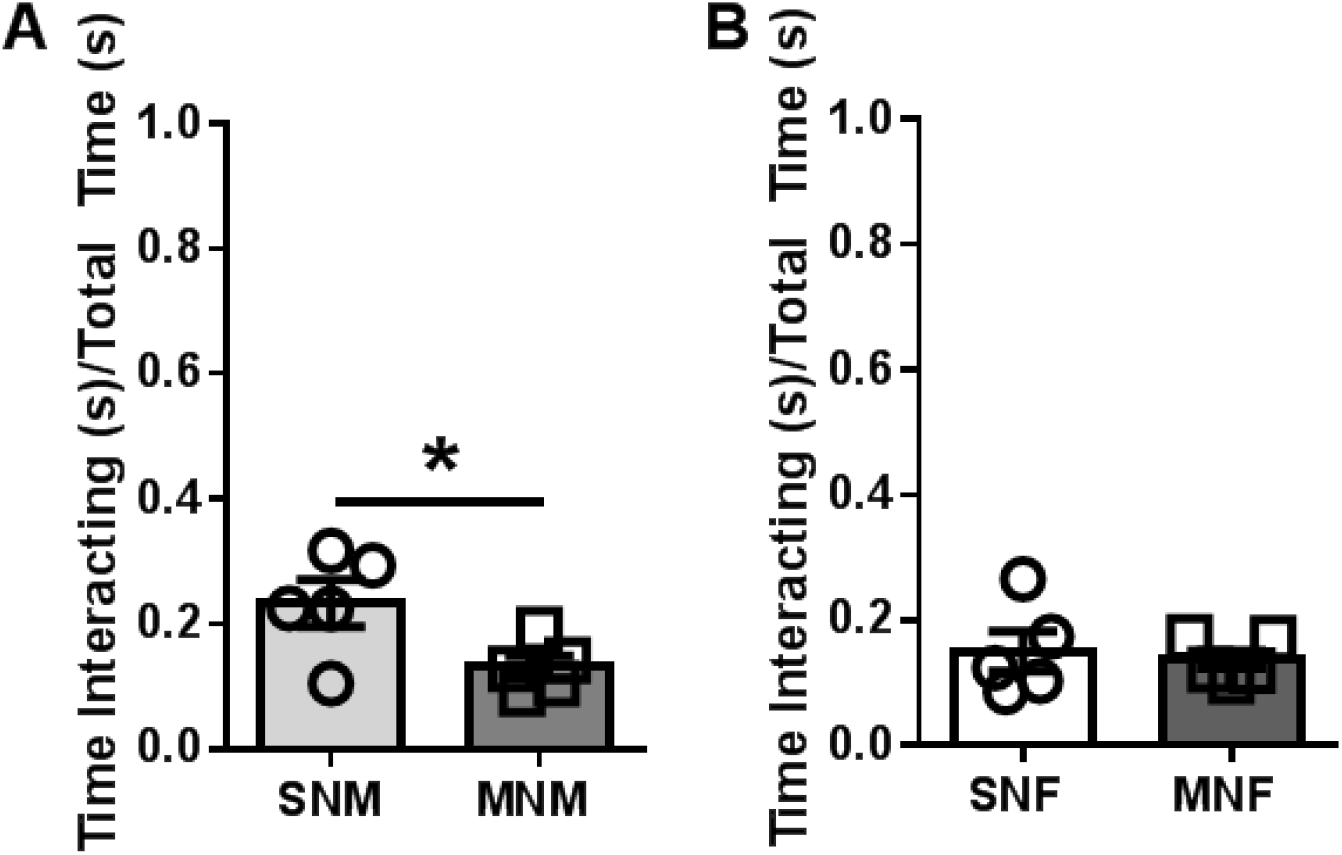
Protracted withdrawal alters social interaction time in male but not female mice. A. Time spent interacting between male experimental mice and age-matched conspecific target mice. B. Time spent interacting between female experimental mice and age-matched conspecific target mice. * p>0.05

### Locomotor sensitization in protracted withdrawal

We next wanted to investigate if we would observe lasting changes in locomotor responsivity to challenge doses of morphine. Based on previous work (Rothwell *et al.*, 2010), we hypothesized that animals with a history of opioid withdrawal would show enhanced locomotion following acute doses of morphine. Six weeks following our morphine withdrawal paradigm, animals were injected with saline, 1 mg/kg, 5 mg/kg, and 10 mg/kg morphine (with at least two days between each injection to allow for total drug clearance) and allowed to explore an open field for a total of 150 minutes. In male mice, SNM and MNM did not show differences in distance traveled following saline **(Fig 7A),** however there was a main effect of withdrawal history on distance traveled following 1 mg/kg morphine (**Fig 7B**, F_(1,18)_=5.987, p<0.05), and significant withdrawal history × time interaction following 5 mg/kg morphine (**Fig 7C**, F_(14,252)_=2.654, p<0.01). At a dose of 10 mg/kg morphine we did not observe any significant differences between our groups in distance traveled in the open field in the male mice (**Fig 7D**). Similarly, in female mice, withdrawal history did not alter the distance traveled in the open field following a dose of saline (**Fig 7E**), 1 mg/kg (**Fig 7F**), or 5 mg/kg morphine (**Fig 7G**). A challenge dose of 10 mg/kg morphine however, produced a significant main effect of prior withdrawal history (**Fig 7H**). These data suggest that both male and female mice that experienced withdrawal have a sensitized locomotor response to subsequent morphine, although to different doses.

**Figure 7.**
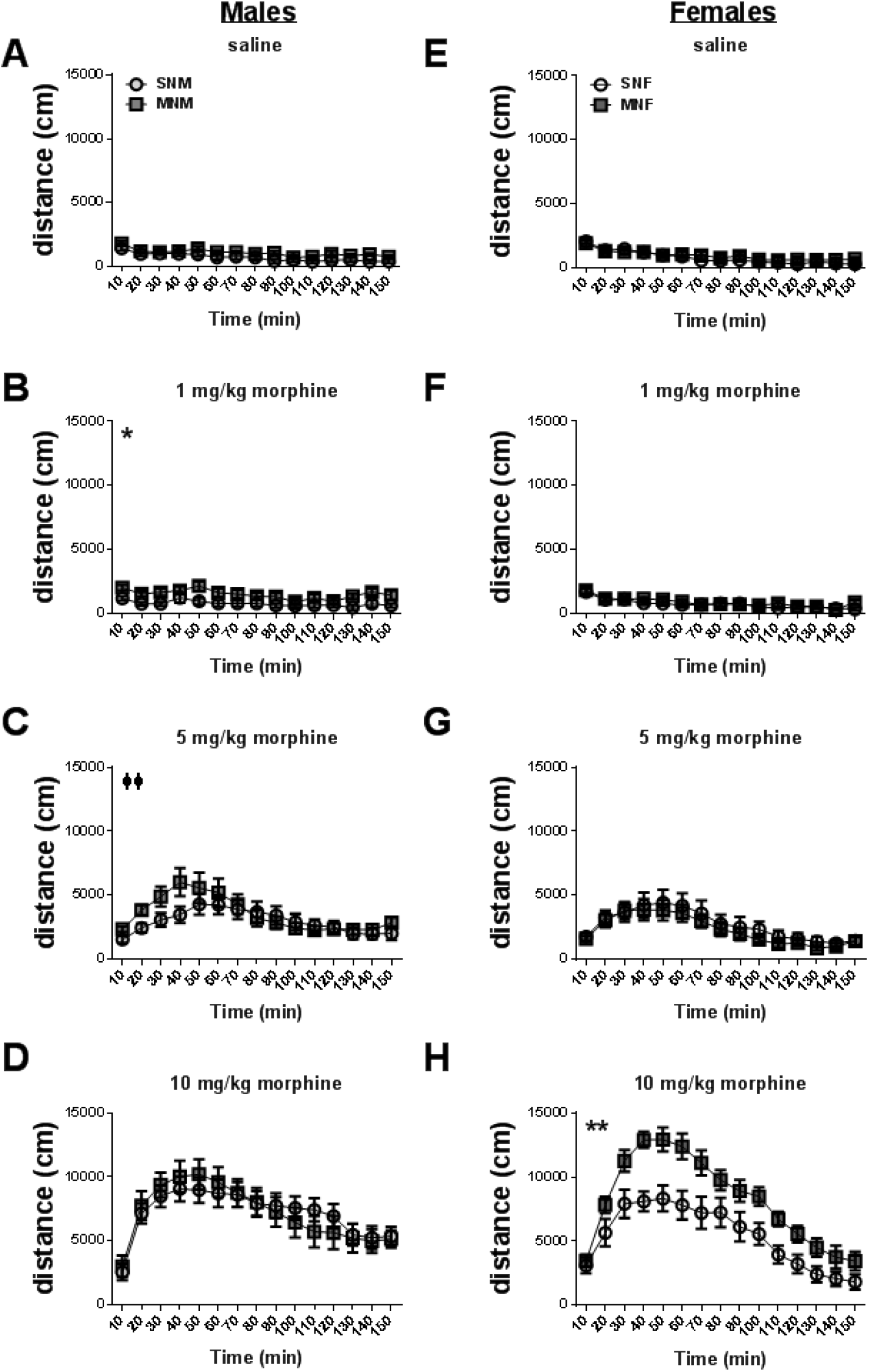
Locomotor sensitization to morphine in male and female mice in protracted withdrawal. Distance traveled in the open field of SN and MN male mice injected subcutaneously (s.c.). A. saline, B. 1 mg/kg morphine, C. 5 mg/kg morphine, D. 10 mg/kg morphine. Distance traveled in the open field of SN and MN female mice injected s.c. E. saline, F. 1 mg/kg morphine, G. 5 mg/kg morphine, H. 10 mg/kg morphine. Main effect of drug treatment: * p<0.05, ** p<0.01. Drug treatment by time interaction: ◻◻ p<0.01

While we did not observe significant differences in the distance traveled following the saline injection (Fig7A,E), we were interested to investigate if the saline injection would act as a mild stress stimulus, thus altering exploration of the open field. We found that male mice that had previously experienced morphine withdrawal spent more time in the center of the open field compared to controls (SNM: 3.4. ± 0.8% vs. MNM: 16.9 + 3.8%, n=10 both groups, p<0.01, **Fig8A**), and had significantly greater number of entries into the center of the open field (SNM: 259.0 ± 82.8 entries vs. MNM: 676.7 ± 90.0 entries, p<0.01, **Fig8C**). In contrast, we did not observe any significant differences between female mice with a history of morphine withdrawal and controls in either center time (SNF: 10.2 ± 2.2% vs. MNF: 10.8 ± 3.7%, n=10 both groups, **Fig8B**), or entries into the center (SNM: 402.2 ± 59.2 entries vs. MNM: 470.5 ± 142.9 entries, p<0.01, **Fig8D**). These data suggest that the saline injection alters exploration in male, but not female mice with a history of morphine withdrawal.

**Figure 8.**
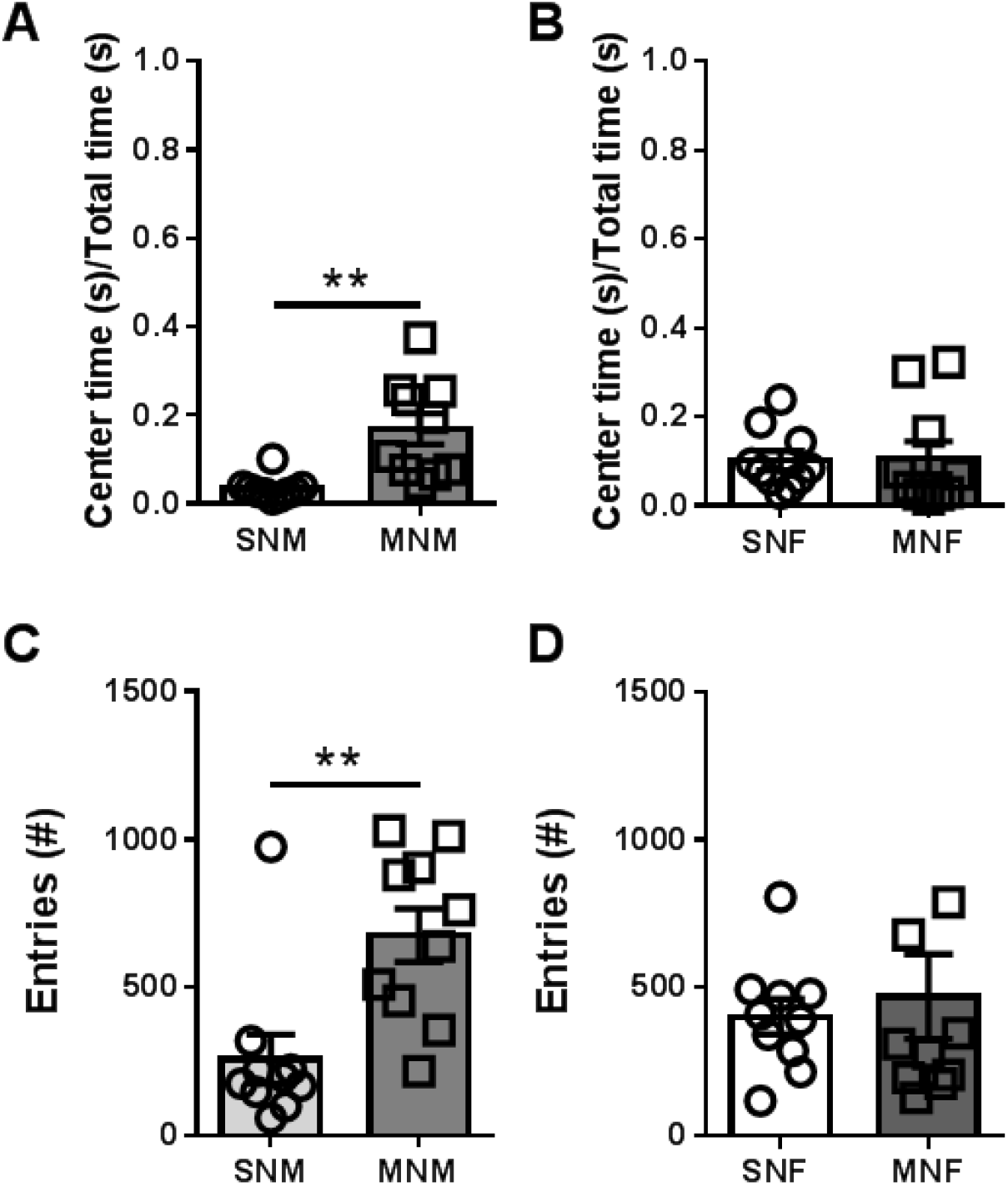
Protracted withdrawal alters stress responsivity in male mice. A. Ratio of center time over total time in the open field in male mice injected with saline. B. Ratio of center time over total time in the open field in female mice injected with saline. C. Number of entries into the center of the open field in male mice injected with saline. D. Number of entries into the center of the open field in male mice injected with saline. * p<0.05, ** p<0.01

## Discussion

In this study we investigated multiple aspects of anxiety-like behavior along with locomotor responses to a morphine challenge in male and female C57BL/6J mice that had previously experienced an acute morphine dependence paradigm. While we found that both male and female mice show altered behavior six weeks following three days of precipitated morphine withdrawal, these behaviors were differentially manifested across both sexes. These data demonstrate that precipitated withdrawal from a low dose of morphine (10 mg/kg), results in long-lasting behavioral plasticity.

### Morphine withdrawal paradigm

It is well established that withdrawal from drugs of abuse is a contributing factor to the development of substance use disorder (Koob & Volkow, 2010). Examining opioid dependence and withdrawal in mouse models, however, presents interesting challenges to biomedical researchers because it is difficult to replicate the means by which humans consume opioids. To attempt to model the physiological and behavioral changes in OUD, many researchers use relatively high doses of morphine, and/or escalating doses to induce both acute and chronic physical dependence (Kest *et al.*, 2001; Goeldner *et al.*, 2011; Valentinova *et al.*, 2019). It has been observed, however, that acute dependence can be induced with a rewarding, albeit lower dose of morphine (10 mg/kg) and precipitated withdrawal with relatively low doses of the opioid receptor antagonist naloxone (1 mg/kg)(Schulteis *et al.*, 1998, 1999; Le Merrer *et al.*, 2012; Neelakantan *et al.*, 2016). Investigating physiological and behavioral changes to lower doses of opioids may be clinically relevant in the current opioid crisis, as 80% of new heroin users report previous prescription opioid use (Cicero & Kuehn, 2014). Furthermore, the intensity of somatic withdrawal symptoms (Fishman, 2008) (MCHUGH 2016) and the distress associated with experiencing withdrawal contributes lack of treatment compliance and opioid misuse (McHugh *et al.*, 2016). Because of these observations in human populations, we chose to use a model where animals receive a rewarding/analgesic dose of morphine, yet experience substantial withdrawal (see **Fig1**), in an attempt to capture the transition between dependence and opioid misuse.

Recently we demonstrated that this paradigm sensitizes the expression of somatic withdrawal behaviors in both male and female C57BL/6J mice, and alters inhibitory transmission in the bed nucleus of the stria terminalis (BNST), a brain region implicated in mediating anxiety-like behavior in both rodents and humans (Cecchi *et al.*, 2002; Mazzone *et al.*, 2016; Buff *et al.*, 2017; Fitzgerald *et al.*, 2018). Previously, we observed sex specific manifestations of withdrawal behavior and subsequent BNST physiology (Luster *et al.*, 2019). In the present study, we show that our 3-day morphine exposure paradigm (see **Fig1)** produces behavioral changes indicative of a protracted withdrawal syndrome in male and female C57BL/6J mice that persists 6 weeks following the last drug treatment. These data suggest that our paradigm captures multiple elements of the human experience.

### Protracted withdrawal alters the pattern of maze exploration

Although physical withdrawal symptoms (vomiting, diarrhea, flu like symptoms, etc.) subside shortly after cessation of opioid use, protracted withdrawal symptoms, primarily presenting as negative emotional valence (e.g. anxiety, depression, social anxiety, etc.) have been well documented in clinical settings (citations). To investigate anxiety-like behaviors at baseline we compared MN male and female mice 6 weeks into protracted withdrawal with SN-treated controls. We used two separate assays that present mice with a conflict between their exploratory nature and desire to remain safe in more enclosed environments: the elevated plus maze (EPM), and the open field (OF). While the canonical interpretation is that mice with enhanced anxiety will spend more time in the closed arms of the EPM and less time in the center of the OF, male C57BL/6J mice are known to respond in opposition. A detailed investigation into stress responsivity from the Holmes Lab demonstrated that in contrast to other strains, C57BL/6J mice spend more time in exposed areas following chronic stress, which they interpreted as an “adaptive coping strategy” (Mozhui *et al.*, 2010). Akin to this finding, male C57BL/6J mice 6 weeks into protracted withdrawal from our opioid paradigm show increased time spent, and entries to the open arms in the EPM (**Fig2**), and increased latency to enter the corner of the OF (**Fig4**). In contrast, female mice 6 weeks following their last drug treatment did not show any differences in the EPM (**Fig3**), and showed reduced time spent in the center, and enhanced time spent in the corners of the OF **(Fig5).** These data suggest that protracted withdrawal behavior is differentially manifested and expressed in males and females, thus the use of various maze/conflict assays will be important in future research, as evidence of protracted withdrawal may be observed in some assays, but not others.

While these measures captured the mice at baseline, we reasoned that these mice might respond differently if presented with a stress challenge. Thus, we analyzed how the mice responded to an acute mild stressor (saline injection) in the OF, which was built into the control of our morphine challenge experiment. Similar to the EPM data, male mice with a history of opioid withdrawal spent significantly more time in, and exhibited increased entries into the center of the OF following this mild stressor, while female mice did not **(Fig8).**

One potential interpretation of the behavior of our male mice following protracted morphine withdrawal is that they are showing behavioral disinhibition, behaving in an impulsive-like manner (e.g. faster entry into to the open arm of an EPM out of impulsivity, rather than a slower latency, which is interpreted canonically as an anxiety-like response). A recent study has suggested that the circuit between the central nucleus of the amygdala and the BNST may regulate impulsive behavior (Kim *et al.*, 2018). Given that we find sexual dimorphism in inhibitory transmission in the BNST following acute morphine withdrawal (Luster *et al.*, 2019), these data suggest a potential mechanism for this long-lasting behavioral plasticity. It will be intriguing to investigate the physiology of these extended amygdala circuits following protracted morphine withdrawal, and the role of these circuits in impulsivity in future studies of protracted withdrawal. A second interpretation could be that the male C57BL/6J mice displaying an active coping strategy versus a passive coping strategy when faced with a stressor (the novel mazes and saline injection). Indeed, others have shown that alcohol withdrawal in C57BL/6J male mice results in an increase in marble burying behavior (Pleil *et al.*, 2015; Rose *et al.*, 2016), thought to be an active coping strategy. In future studies, we will investigate defensive burying behavior following protracted withdrawal from this morphine paradigm.

In contrast to the MN male mice, MN female C57BL/6J mice spent less time in the center and more time in the corners of the OF as compared to their SN controls, which we interpret as an increase in anxiety-like behavior. Interestingly, however examining the time course it is evident that this behavior did not manifest in the first 15 minutes of the experiment, but rather appeared in the middle of the testing period. This suggests that we may have not observed differences in the EPM with in female mice because of the short duration of the test (5 minutes). These data show that female C57BL/6J mice may (1) exhibit behavior completely opposite of males in protracted morphine withdrawal and (2) that anxiety-like behavior in female C57BL/6J mice may not be readily apparent until later in the assay.

### Social interaction deficits in protracted morphine withdrawal

Deficits in social interaction behavior are associated with drug withdrawal and anxiety-like behavior (Marcinkiewcz *et al.*, 2015; Valentinova *et al.*, 2019). Previous research has demonstrated that morphine dependence induced by escalating doses (20-100 mg/kg) across 6 days in male mice results in a deficit in social interaction in 4 weeks into protracted withdrawal (Goeldner *et al.*, 2011). To observe whether our model would produce similar deficits, we conducted a social interaction test in MN- and SN-treated mice 6 weeks into protracted withdrawal. Similar to Goeldner *et al.*, male C57BL/6J mice in protracted withdrawal had a significant deficit in social interaction with a naïve age-matched conspecific. We did not observe any differences, however, in the female animals. In light of our results in the OF, where differences were not observed until at least 15-30 minutes into the session, future studies examining social interaction in female animals following protracted withdrawal from our paradigm will use longer periods of interaction which may reveal a time dependent effect.

### Protracted morphine withdrawal enhances the locomotor effect of a challenge dose of morphine

One of the most well studied aspects of repeated drug administration in rodent models is a sensitization of the psychomotor-activating properties of drugs of abuse (Berridge & Robinson, 2016). A previous study using a similar dosing regimen to our paradigm by (Rothwell *et al.*, 2010) demonstrated that exacerbating withdrawal with repeated naloxone administration enhanced the observed locomotor sensitization to lower doses of morphine a week into protracted withdrawal from the last morphine/naloxone pairing. Because our data suggested that there was long lasting behavioral adaptation in anxiety-like measures, we also wanted to determine if there would be an enhanced response to challenge doses of morphine during protracted withdrawal. We found that both male and female MN-treated animals exhibited dose dependent differences in their locomotor response to subsequent morphine challenge. Male animals showed enhanced locomotor effects following both 1 and 5 mg/kg morphine doses, while the effect in females was not apparent until 10 mg/kg morphine **(Fig. 7).** Sex differences in analgesic responses to opioids have been observed in both humans and mice, wherein females are less sensitive to the pain-relieving properties of these drugs (Kest *et al.*, 2000). However, females also show equal or enhanced responses to the rewarding properties of opioids (Le Merrer *et al.*, 2012; Neelakantan *et al.*, 2016; Pleil & Skelly, 2018), suggesting a dissociation between these two interoceptive states in response to opioids. Because of the association between locomotor sensitization and compulsive drug seeking and taking (Robinson & Berridge, 2003; Vezina, 2004), the lasting effects we observe here in protracted withdrawal may be highly significant for potential propensity to relapse.

In this study, we have examined long-lasting changes in behavioral plasticity following a 3-day paradigm of morphine exposure and precipitated withdrawal. The goal of this study was to determine whether our model would produce long-lasting behavioral changes after repeated naloxone-precipitated withdrawal from morphine in male and female mice. Here we report that 6 weeks following morphine exposure and withdrawal we find significant increases in anxiety-like behavior in male and female morphine-treated C57BL/6J mice, however the manifestation of these behaviors is sex specific. Similarly, we observe dose dependent enhanced locomotor responsivity to challenge doses of morphine that differ between male and female mice that were previously treated with morphine and naloxone, as compared to their controls. Taken together, these studies provide support for the clinical relevance of our 3-day withdrawal paradigm (with a relatively low dose of morphine) as a model to investigate enduring behavioral adaptation in both male and female animals. These results are congruent with what has been observed in other models using higher doses of morphine, and shorter periods of protracted withdrawal(Rothwell *et al.*, 2009, 2010; Goeldner *et al.*, 2011; Valentinova *et al.*, 2019). By understanding the nuance, similarities, and differences in male and female animals on the behavioral level (Shansky, 2019) we are better equipped to discover neural mechanisms underlying how withdrawal experience leads to vulnerabilities for substance use disorders.

## Acknowledgements

This work was supported by: K01AA023555 (Z.A.M.), T32 AA007573 (M.E.F, E.S.C., K.T.S.), T32 MH093315 (B.R.L.), R25 GM089569 (P.J.P.). We thank Dr. Thomas Kash for comments on a previous version of this manuscript.

